# Disturbed ATP and AMPK homeostasis in an Ank_F377del_ mouse model for craniometaphyseal dysplasia

**DOI:** 10.64898/2026.04.15.717889

**Authors:** Ayano Hatori, Shyam Kishor Sah, Koen van de Wetering, Ernst J Reichenberger, I-Ping Chen

## Abstract

Craniometaphyseal dysplasia (CMD) is a rare genetic disorder characterized by hyperostosis of craniofacial bones and flared metaphyses of long bones. Mutations in ANKH (mouse orthologue ANK), a transmembrane protein mediating ATP and citrate efflux, cause the autosomal dominant form of CMD. How ANK mutations in CMD affect ATP/citrate homeostasis and downstream targets remains unknown. We determined that cellular ATP export, intracellular ATP levels, and plasma citric acid were significantly reduced in ANK_F377del_ knock-in (*Ank^KI/KI^*) mice. Enrichment and pathway analyses of the plasma metabolome suggested the involvement of the citric acid cycle. It is known that AMPK is phosphorylated and activated when ATP is low. Phospho-AMPK was significantly upregulated in fusing *Ank^KI/KI^* osteoclasts, major contributors to CMD. AMPK inhibitor treatment only during the fusion stage of osteoclasts significantly restored dysfunctional *Ank^KI/KI^* osteoclasts, partly by modulating actin structures. Systemic administration of the AMPK inhibitor SBI-0206965 improved the positioning of cervical loops of incisors but failed to correct other skeletal abnormalities in *Ank^KI/KI^* mice. Limitations of systemic administration of SBI-0206965 include its off-target effects on other cell types and the inability to inhibit AMPK only on fusing osteoclasts. Nonetheless, this proof-of-principle study reveals an important role of the ATP-AMPK axis in CMD pathogenesis.

**Take-home message:** Suppression of increased activation of AMPK restores the function of osteoclasts, suggesting that abnormal energy metabolism is an integral component of the disease phenotype in CMD.

## INTRODUCTION

CMD is characterized by hyperostosis of craniofacial bones and flared metaphyses of long bones.^(1)^ Individuals with CMD often suffer from deafness, blindness, facial palsy, and in severe instances, patients may have reduced life expectancy due to bony compression of the spinal cord.^(2, 3)^ CMD can be inherited as an autosomal dominant (AD, OMIM 605145) trait with mutations identified in the progressive ankylosis (*ANKH*) gene and in an autosomal recessive form (AR, OMIM 121014) carrying a mutation in the gap junction protein alpha 1 gene (*GJA1*).^(4–6)^ Diagnosis of CMD is based on clinical examination, radiographic findings, and identification of genetic mutations. Clinical care for CMD includes close monitoring of craniofacial bone thickening with foraminal encroachment, management of vision or hearing loss, and repetitive decompression surgeries as needed. To date, there is no effective treatment for CMD.

We have generated a knock-in (KI) mouse model (*Ank^KI/KI^* mice) carrying the AD CMD mutation ANK_F377del_. *Ank^KI/KI^* mice replicate many features of craniofacial and long bones of human CMD.^(7)^ Mice with ablation of the ANK protein (*Ank^ank/ank^* and *Ank^null/null^*) share some phenotypes with our *Ank^KI/KI^* mice, but do not fully phenocopy human CMD. Some of the most characteristic features of CMD such as club-shaped femurs, severe narrowing of cranial foramina, and massive jawbones are only found in the *Ank^KI/KI^* mice.^(7–9)^ *Ank^+/KI^* mice initially appear phenotypically normal but develop intermediate CMD-like features after approximately one year. We previously showed that CMD-mutant osteoclasts (OCs) are major contributors to the CMD phenotype.^(10)^ Mutant ANK_F377del_/ANKH_F377del_ protein results in significantly reduced OC formation and decreased bone resorption.^(7, 10, 11)^ *Ank^KI/KI^*OCs are slower to migrate and smaller in size with disrupted actin rings.^(10)^ Although *in vitro* osteoblast (OB) differentiation is remarkably impaired, bone formation rate and mineral apposition rate are not significantly affected in *Ank^KI/KI^* mice.^(7),(10)^

The ANKH (mouse ortholog ANK) transmembrane protein has previously been described as a transporter for pyrophosphate (PPi).^(12)^ The presence of PPi in non-bone tissues is critical to prevent ectopic calcification.^(13)^ Recent studies have shown that ANKH exports small molecules including citrate and adenosine triphosphates (ATP), instead of PPi.^(14–16)^ Decreased extracellular PPi is a result of reduced ATP export in cells with loss-of-function ANK.^(14–16)^ We and others have shown that CMD-mutant ANK causes decreased extracellular PPi levels in long bones.^(17, 18)^ How the ATP/citrate homeostasis and downstream targets are affected remains unknown.

AMPK, a heterotrimeric protein consisting of a catalytic α and regulatory β and γ subunits, is a key energy sensor to tightly maintain cellular and systemic energy homeostasis.^(19)^ AMPK is activated via the phosphorylation of Thr-172 within the α subunit to reduce ATP consumption and increase ATP production during glucose deprivation or when ATP is low.^(20, 21)^ AMPK also regulates bone remodeling by acting on OCs and OBs. The deletion of AMPK catalytic α subunits in *Prkaa1^−/−^* and *Prkaa2^−/−^* mice results in reduced bone mass due to increased OC formation and cell fusion suggesting that AMPK is a negative regulator of osteoclastogenesis.^(22)^ The effects of AMPK on osteoblasts are less consistent. *Prkaa1^−/−^*mice have increased OB-mediated bone formation while *Prkaa2^−/−^*mice display no detectable changes in osteoblast activity.^(22)^

In the present study, for the first time, we show a disturbed ATP and citrate metabolism in *Ank^KI/KI^* mice. We linked impaired osteoclastogenesis to increased activation of AMPK in *Ank^KI/KI^* fusing OCs *in vitro*. These novel findings suggest a previously unrecognized association between energy metabolism and CMD pathogenesis.

## MATERIALS AND METHODS

### Mouse BMM cultures

*Ank^KI/KI^* mice were first generated in 129S1/SvlmJ background and backcrossed with C57BL/6J mice for 9 generations.^(7)^ Mice were housed in an AAALAC-accredited facility under veterinary supervision. BMM cultures were prepared from tibia and femurs of 6- to 9-week-old *Ank^+/+^* and *Ank^KI/KI^* mice as previously described.^(10)^ Bone marrow was flushed out and cultured in α-MEM medium containing 10% FBS (Atlanta Biologicals) and penicillin-streptomycin (Thermo Fisher Scientific). Non-adherent cells were collected and purified by Ficoll separation (StemCell Technologies) after 18-24 hours. BMMs were seeded (cell density: 3×10^5^/well on 6-well plates, 5000/well on 96-well plates, 10,000 per bone chip) and grown in α-MEM medium supplemented with 30 ng/mL M-CSF (PeproTech) for the first 2 days to enrich OC progenitors followed by the treatment of 30 ng/mL M-CSF and 30 ng/mL RANKL (PeproTech) to promote OC differentiation. Media were changed every two days.

BMMs cultured on 96-well culture plates or bone chips were treated with and without AMPK inhibitors compound C or SBI-0206965 (Selleck Chemicals) at concentrations of 0.5 μM. Cells were first cultured in M-CSF for 2 days followed by M-CSF and RANKL. Cells cultured in M-CSF and RANKL were treated with inhibitors during the fusion stage (days 0-2), the fusion to maturation stage (days 2-5) or by continuous treatment (days 0-5). All cells were terminated and analyzed on day 5 of M-CSF and RANKL treatment.

### ATP and ADP assays in BMM cultures

BMMs cultured on 100-mm-dishes were treated with M-CSF for 2 days to enrich OC progenitors, followed by treatment with M-CSF and RANKL for 1 day. Following overnight incubation with M-CSF and RANKL, BMMs were trypsinized and replated at a density of 5,000 cells per well in opaque-walled 96-well plates containing 100 μL of complete medium. The cells were incubated overnight at 37 ℃ in a humidified incubator with 5% CO_2_.

ATP efflux and Intracellular ATP assays of BMMs were performed using RealTime-Glo Extracellular ATP Assay (Promega) and CellTiter-Glo 2.0 Assay (Promega), respectively, according to the manufacturer’s protocols. For extracellular ATP assays, 1250 nM mitoxantrone, an ATP release inducer, was added along with RealTime-Glo extracellular ATP assay substrate. Luminescence was measured from 96-well white opaque plates (Thermo Fisher Scientific) using a SpectraMax microplate reader (Molecular Devices). Luminescence was measured kinetically every 30 minutes for up to 8 hours. ATP efflux and intracellular ATP assays were performed in 4-6 wells for three experiments.

ADP assay was performed using the ADP assay (fluorometric) kit (Abcam) according to the manufacturer’s protocols. BMMs cultured under the same condition as for ATP assays described above were trypsinized and counted. 4 x 10^6^ cells were used for each assay. Briefly, cells were washed with cold PBS, resuspended in ADP assay buffer, homogenized by pipetting up and down, and incubated on ice for 10 minutes. The supernatant, after spinning down, was collected and transferred to the wells of black 96-well plates (Thermo Fisher Scientific). A standard curve was prepared with duplicate readings. After adding the reaction mix, the plate was incubated at room temperature in the dark for 30 minutes. Fluorescence was measured at an Ex/Em of 535/587 nm using a microplate reader (Molecular Devices). The assay was performed for three experiments.

### Untargeted metabolomics study

We collected and subjected plasma samples from 12-week-old male and female *Ank^+/+^* and *Ank^KI/KI^* mice (male *Ank^+/+^*: n=10, male *Ank^KI/KI^*: n=8, female *Ank^+/+^*: n=8, female *Ank^KI/KI^*: n=5) for untargeted metabolomics analysis at the UConn Proteomics & Metabolomics Facility. Metabolites were isolated using methanol liquid phase extraction (methanol/plasma 4:1 ratio). Methanol extracts were diluted in LCMS grade water (Fisher Scientific), frozen at −80 °C, and lyophilized (Labconco). The dried extracts were reconstituted in 0.1% formic acid in water. The reconstitution volume was adjusted to normalize the sample concentration at 10 mg/mL. The samples were then centrifuged at 12,500 rpm for 10 min, and the supernatant was transferred to an LCMS vial (Waters Corporation). The samples were analyzed via tandem mass spectrometry on a Xevo G2-XS UPLC-MS/MS instrument (Waters Corporation) operated in both positive and negative electrospray ionization modes. Chromatographic separations were performed using an Acquity UPLC HSS T3 column (150 x 2.1 mm, 1.8 um particle size) (Waters Corporation). Water and acetonitrile, both containing 0.1% formic acid were used as solvents A and B respectively; and water and acetonitrile, both with 10 mM triethylammonium acetate were used as solvents C and D respectively. The injection volume was 10 μL. The mobile phase gradient at 0.3 mL/min was maintained as follows, 0 - 0.5 min: 5% B/D; 8.5 – 9 min: 98% B/D; and 9.25 – 12.5 min: 5% B/D. The mass range for MS and MS2 acquisitions were 110 to 2000 Da and 50 to 2000 Da, respectively. The UPLC-MS/MS raw data files were analyzed via Progenesis QI software (v 2.4, Waters Corporation) to perform retention time alignment, peak picking, cross-sample normalization, and adduct deconvolution. Compound identification was performed using the METLIN MS/MS library. Search parameters were set to include 10 ppm precursor tolerance and 20 ppm fragment tolerance. Data processing and model fitting were performed by the UConn Statistical Consulting Services. MetaboAnalyst 6.0 (metaboanalyst.ca) was used for data analysis to identify significant metabolites, enrichment analysis, and pathway analysis with a fold-change ≥ 2, raw p value < 0.05, and −log_10_(p) >1.5. Metabolomics data are included in Supporting information.

### Immunoblotting

Whole cell lysate was prepared in RIPA buffer (150 mM NaCl, 1% NP-40, 0.5% sodium deoxycholate, 0.1% SDS, 50mM Tris (pH 7.4), 1x phosphatase and protease inhibitor cocktail). Protein was quantified using a Pierce BCA protein assay kit (Thermo Fisher Scientific). Equal amounts of protein were subjected to SDS-polyacrylamide gel electrophoresis and transferred to PVDF membranes using a tank transfer system (Bio-Rad Laboratories). Membranes were blocked in 3% BSA for one hour at room temperature and incubated with specific antibodies p-AMPK_Thr172_ 1:1000 (Cell Signaling Technology); AMPK 1:1000 (Cell Signaling Technology); actin 1:1000 (Santa Cruz Biotechnology) at 4°C overnight. The next day, membranes were washed and incubated with the corresponding secondary antibody for 1 hour at room temperature. Signals were visualized using Radiance Q chemiluminescent ECL substrate (Azure Biosystems). Quantification was performed by densitometry analysis using ImageJ software (National Institutes of Health, NIH). Whole blots of presented figures are shown in Supplementary Figure S1.

### Analyses of BMM cultures

For tartrate resistant acid phosphatase (TRAP) staining, cells were fixed with 2.5% glutaraldehyde at room temperature for 30 minutes. TRAP staining was performed using an acid phosphatase leukocyte kit (Sigma-Aldrich) following manufacturer’s instruction. Images were taken by a Rebel microscope (Echo). TRAP-positive cells with 3 or more nuclei were counted as OCs.

We examined actin rings of OCs by rhodamine-phalloidin staining for 25 minutes in the dark (1:40 dilution in PBS; Invitrogen-Molecular Probes). Nuclei were stained with Hoechst 33342 dye (Molecular Probes; 1:1000 in PBS). Images were taken by a Z1 Observer microscope (Zeiss). Surface area of OC and size of actin ring were quantified by Image J software (NIH).

For bone resorption pit assays, BMM cultures on bone chips were terminated at day 10. Bone chips were air dried and imaged using a TM-1000 tabletop scanning electron microscope (Hitachi). To analyze the resorption pits on bone chips, images from bone chips were taken and the percent resorption was calculated as the ratio of resorbed area to total area using Image J software (NIH).

We analyzed acidic vesicles by quinacrine dihydrochloride (QA, 0.3 μM) in *Ank^+/+^* and *Ank^KI/KI^* mature OCs as previously described.^(23)^ QA stock solution was freshly made by dissolving QA (Sigma-Aldrich) in PBS at a concentration of 0.15 μg/mL. The stock solutions were further diluted in culture media. Briefly, cells were washed three times with live cell imaging solution (Invitrogen) and incubated with QA dye for 10 minutes prior to imaging. QA exclusively stains acidic vesicles exclusively under these treatment conditions. Images were taken using an Observer Z1 microscope (Zeiss). The intensity of QA (green) fluorescence was quantified by Image J software (NIH).

### Systemic administration of AMPK inhibitor SBI-0206965

SBI-0206965 (2 mg/mL; MedChemExpress) was prepared in a solvent containing 10% DMSO, 40% PEG300, 5% Tween-80, and 45% saline according to manufacturer’s instructions. *Ank^+/+^* and *Ank^KI/KI^* male and female mice (n = 3-6 per group) were intraperitoneally injected with vehicle (solvent described above) and SBI-0206965 (10 μg/g) weekly starting from 1 to 8 weeks of age.

### Skeletal analysis

Skulls, mandibles, and femurs from 8-week-old *Ank^+/+^* (n=3) and *Ank^KI/KI^* female mice dosed with vehicle (n=6) or SBI-0206965 (n=6) were dissected and fixed in 10% formalin and subjected to radiographic imaging (KUBTEC Scientific) and microcomputed tomography (μCT) (ScanCo Medical AG) in the MicroCT facility at UConn Health. μCT analysis was performed as described previously.^(7)^ In brief, total volume (TV) and bone volume (BV) of mandibles were determined by measuring vertical sections from mesially of 1^st^ molars to distally of 2^nd^ molars. Trabecular measurements were obtained in the metaphyses and cortical parameters were quantified from the diaphyseal region of femurs. Measurements of incisor cervical loops positioning, the width, length, and surface area of foramina magna, and the space of nasal cavity and cranial foramina were determined using Image J software (NIH) (Supplementary Figure S2).

### Statistical analysis

The data are presented as means with standard deviations. Statistical analysis was performed by Student’s *t*-test, one-way or two-way ANOVA followed by multiple comparison tests as indicated, using Prism 5 software (GraphPad Software).

## RESULTS

### Extracellular and intracellular ATP levels are decreased in Ank^KI/KI^ mice

We first examined whether cellular ATP efflux is affected in bone marrow-derived macrophage (BMM) cultures from *Ank^KI/KI^* mice. A luciferase-based real-time ATP efflux assay showed that *Ank^+/+^* BMMs have robust ATP efflux, whereas ATP release from *Ank^KI/KI^* BMMs was significantly reduced (Figure 1A). We suspected that ATP may be accumulated inside cells since the ATP efflux was defective in *Ank^KI/KI^* BMMs. Unexpectedly, intracellular ATP levels, measured by a luciferase reaction generating a luminescent signal that is proportional to the amount of ATP present in cell lysis, were significantly reduced in *Ank^KI/KI^* BMMs compared to *Ank^+/+^* BMMs (Figure 1B). Interestingly, we also detected that intracellular ADP levels were significantly decreased in *Ank^KI/KI^*BMMs by an ADP assay that converts ADP to ATP and pyruvate, which was then fluorometrically quantified (Figure 1C). These data suggest that CMD-mutant ANK results in disturbed ATP and ADP levels in *Ank^KI/KI^* BMMs.

**Figure 1:**
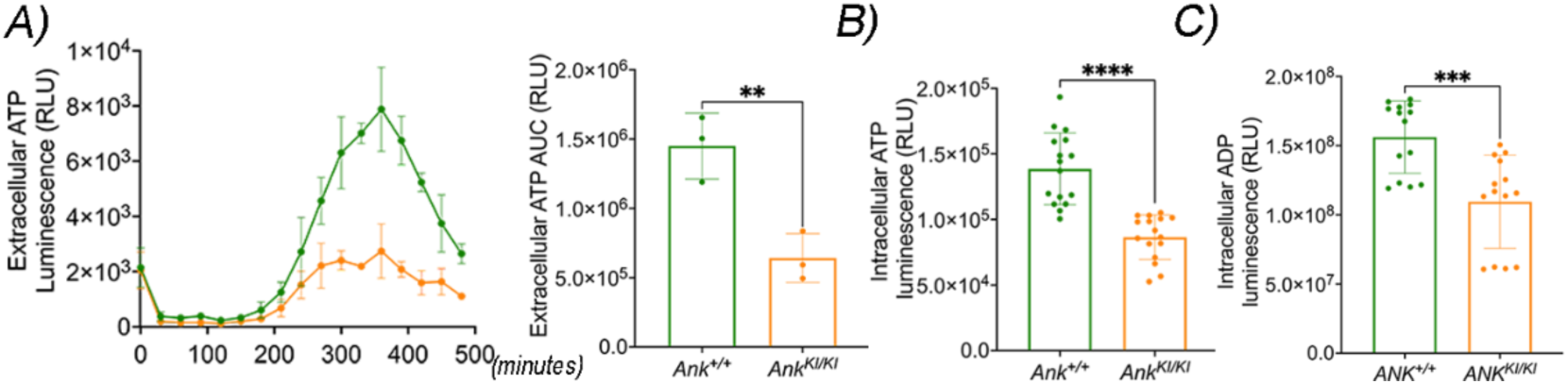
Decreased **A**) release of ATP to extracellular space shown by real-time luminescence signals measured every 30 minutes for 8 hours (left panel) and statistics of extracellular ATP area under the curve (AUC) from three experiments (right panel); **B**) intracellular ATP and **C**) ADP levels in BMM cultures from *Ank^KI/KI^* mice by Student’s *t*-test (\*\**p*<0.01; \*\*\*\**p*<0.001).

### Metabolomic study reveals reduced citric acid in Ank^KI/KI^ plasma

ANK/ANKH has been reported to export small molecules, including citrate.^(14, 24)^ To understand the CMD mutational effects on other metabolites, we performed a plasma metabolome study of *Ank^+/+^*and *Ank^KI/KI^* mice. There were 11,148 compounds detected by positive mode and 300 by negative mode. We merged positive and negative modes and analyzed combined male and female data as shown in Figure 2. Male and female data analyzed separately are shown in Supplementary Figures S3 & S4. We used MetaboAnalyst 6.0 to analyze a total of 451 metabolites that have annotations in the Human Metabolome Database (HMDB). Partial Least Squares Discriminant Analysis (PLS-DA) showed a distinct separation between the *Ank^+/+^* and *Ank^KI/KI^* data (Figure 2A). A variable importance plot (VIP) using comp.1 (T score) listed the top 15 features with most of them being downregulated in *Ank^KI/KI^* mice (Figure 2B). A volcano plot, applying log_10_(p) > 1.5, showed 15 significantly downregulated metabolites, including citric acid and N-acetylneuraminic acid, and 2 upregulated metabolites in *Ank^KI/KI^* mice (Figure 2C). We performed an enrichment assay to link the top 25 identified metabolites with biologically meaningful pathways, conditions or diseases. Our data showed enriched metabolites involved hyperoxalemia and the 2-ketoglutarate dehydrogenase complex (Figure 2D). Pathway analysis indicated that the citric acid cycle was commonly affected in both sexes (Figure 2E, Supplementary Figures S3 & S4). Our data further demonstrated that citric acid and N-acetylneuraminic acid were significantly decreased in both, male and female *Ank^KI/KI^* mice (Figure 2F). Multiple di-, tri-, and tetrapeptides were significantly up- or downregulated in *Ank^KI/KI^* mice although the roles of these metabolites in CMD remains unknown (data not shown).

**Figure 2.**
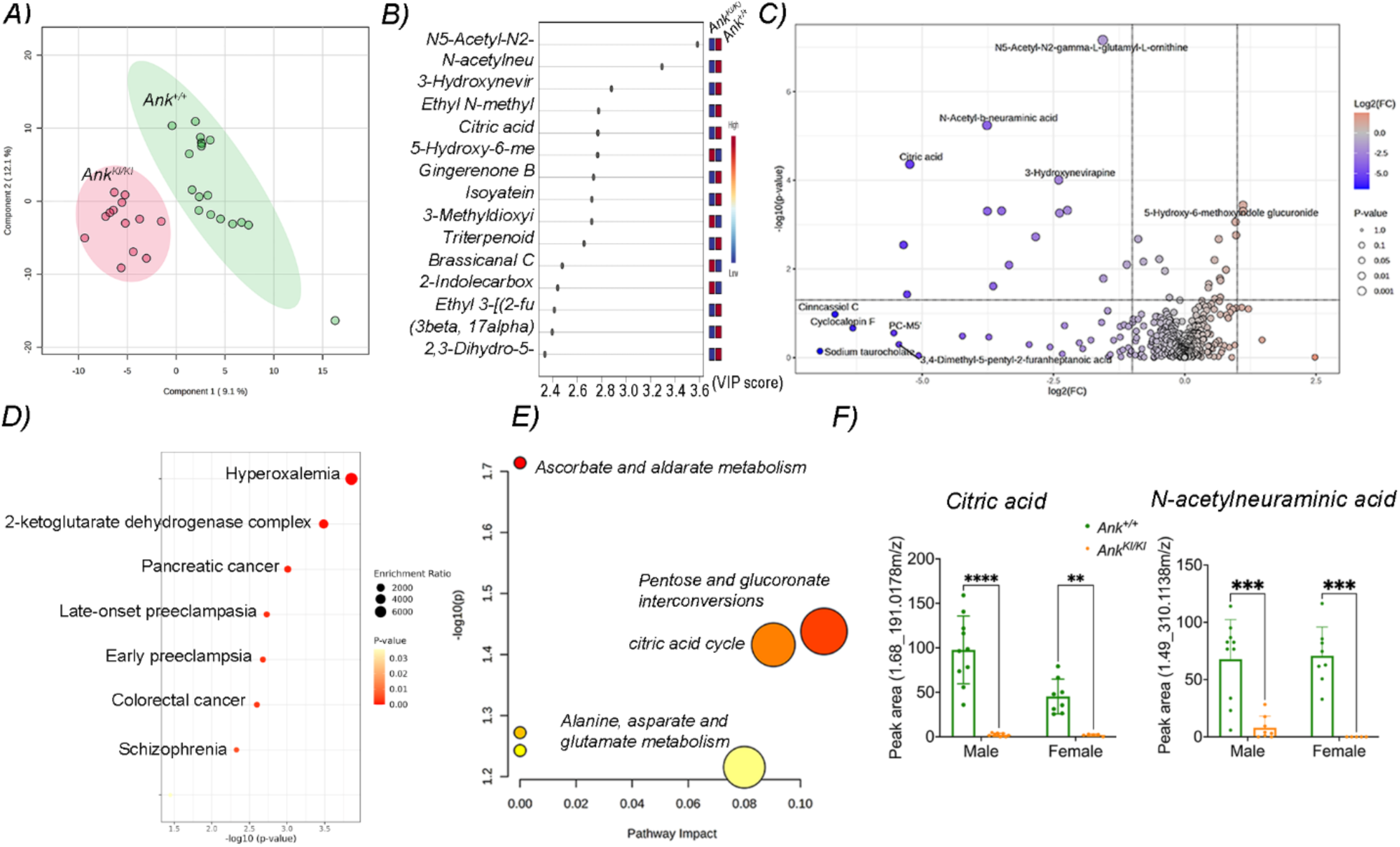
Comparison of plasma metabolome in *Ank^+/+^* and *Ank^KI/KI^* mice. **A**) PLS-DA of combined male and female data and **B**) VIP scores of top 15 differentiated metabolites; **C**) Volcano plot showing significantly upregulated (orange, 2) and downregulated (blue, 15) metabolites (p<0.05, fold-change > 2); **D**) The enrichment assay showing enrichment ratio with *p*-value of upregulated and downregulated top 25 metabolites; **E**) Metabolic pathway analysis showing pathway impact and *p*-value of significantly upregulated and downregulated metabolites; **F**) citric acid and N-acetylneuraminic acid are significantly reduced in both male and female *Ank^KI/KI^* mice. Statistical analysis was performed by two-way ANOVA followed by Sidak multiple comparison test. (\*\**p*<0.01, \*\*\**p*<0.0005, \*\*\*\**p*<0.0001; male *Ank^+/+^* n=10, male *Ank^KI/KI^* n=8, female *Ank^+/+^* n=8, female *Ank^KI/KI^* n=5). Full name of metabolites listed in **B**) are N5-Acetyl-N2-gamma-L-glutamyl-L-ornithine (N5-Acetyl-N2-g); N-Acetylneuraminic acid (N-Acetyl-b-neu); 3-Hydroxynevirapine (3-Hydroxynevir); Ethyl N-methylanthranilate (Ethyl N-Methyl); 5-Hydroxy-6-methoxyindole glucuronide (5-Hydroxy-6-me); 3-Methyldioxyindole (3-Methyldioxyi); 2-Indolecarboxylic acid (2-Indolecarbox); Ethyl 3-[(2-furanylmethyl)thio]propanoate (Ethyl 3-[(2-fu); (3beta, 17alpha, 23S)-17,23-Epoxy-3,29-dihydroxy-27-norlanosta-7,9(11)-diene-15,24-dione (3beta,17alpha); 2,3-Dihydro-5-(3-hydroxypropanoyl)-1H-pyrrolizine (2,3-Dihydro-5).

### Increased activation/phosphorylation of AMPK during Ank^KI/KI^ osteoclastogenesis

When ATP levels are low in cells, AMPK is activated to reduce ATP consumption and increase ATP production.^(25)^ We next examined AMPK phosphorylation during osteoclastogenesis in *Ank^+/+^* and *Ank^KI/KI^* OCs. Cell lysates collected from *Ank^+/+^* and *Ank^KI/KI^* OCs treated with M-CSF and RANKL for 1, 2, and 3 days were subjected to immunoblotting with p-AMPK and total AMPK antibodies. Abundant total AMPK was expressed throughout osteoclastogenesis while p-AMPK increased sharply on day 2 and dropped on day 3 in *Ank^+/+^* and *Ank^KI/KI^* lysates. More importantly, *Ank^KI/KI^* OCs exhibited significantly increased p-AMPK levels at day 2, the fusion stage, compared to *Ank^+/+^* OCs (Figure 3).

**Figure 3.**
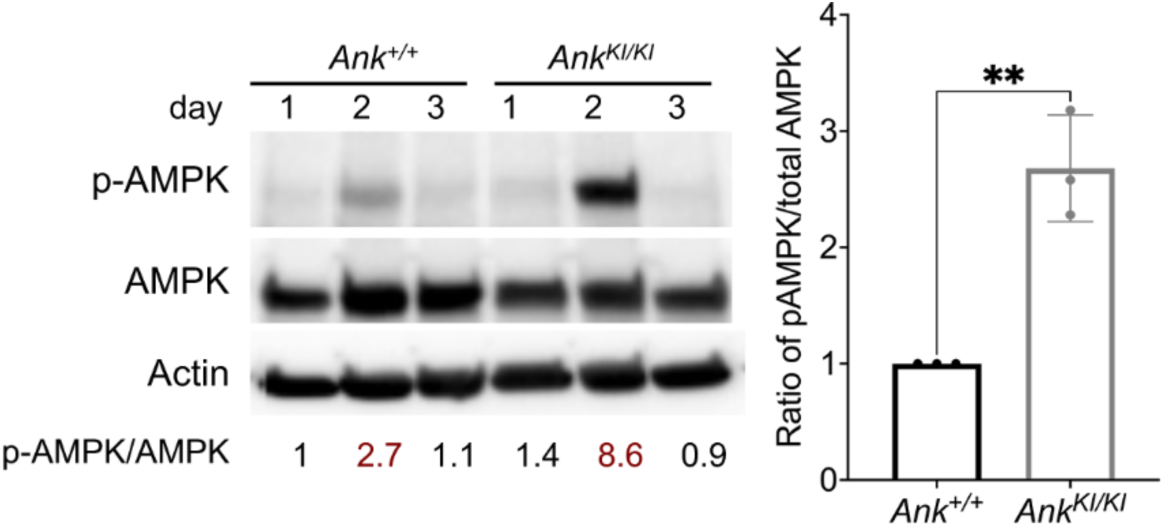
Enhanced AMPK activation during *Ank^KI/KI^* osteoclastogenesis. Increased p-AMPK protein expression in *Ank^KI/KI^* BMMs. AMPK and p-AMPK immunoblotting of lysate from *Ank^+/+^*and *Ank^KI/KI^* OCs cultured in M-CSF for 2 days followed by M-CSF and RANKL for 1, 2, or 3 days. Numbers show ratios of p-AMPK/AMPK normalized to the ratio of *Ank^+/+^* BMMs at day 1. Actin served as loading control. Histogram shows densitometric analysis of p-AMPK/AMPK ratio from biological triplicates of *Ank^+/+^* and *Ank^KI/KI^* OCs at the fusion stage (day 2 in M-CSF and RANKL). Statistics was performed by Student’s *t*-test (\*\**p*<0.01).

### AMPK inhibitor enhances Ank^KI/KI^ osteoclastogenesis in vitro

To study the impact of increased p-AMPK on *Ank^KI/KI^* osteoclastogenesis, we treated BMMs with the AMPK inhibitors compound C or SBI-0206965 at concentrations of 0.5 μM. Compound C is a commonly used AMPK inhibitor but also inhibits multiple other kinases with similar or greater potency.^(26)^ SBI-0206965 selectively inhibits AMPK with 40-fold greater potency than compound C.^(27)^ Cells were treated with AMPK inhibitors during their fusion stage (days 0-2 of M-CSF and RANKL treatment), fusion to maturation stage (days 2-5) or as continuous treatment (days 0-5) (Figure 4A). Analyses of BMM cultures were performed at the end of 5 days of M-CSF and RANKL treatment with or without inhibitors.

**Figure 4.**
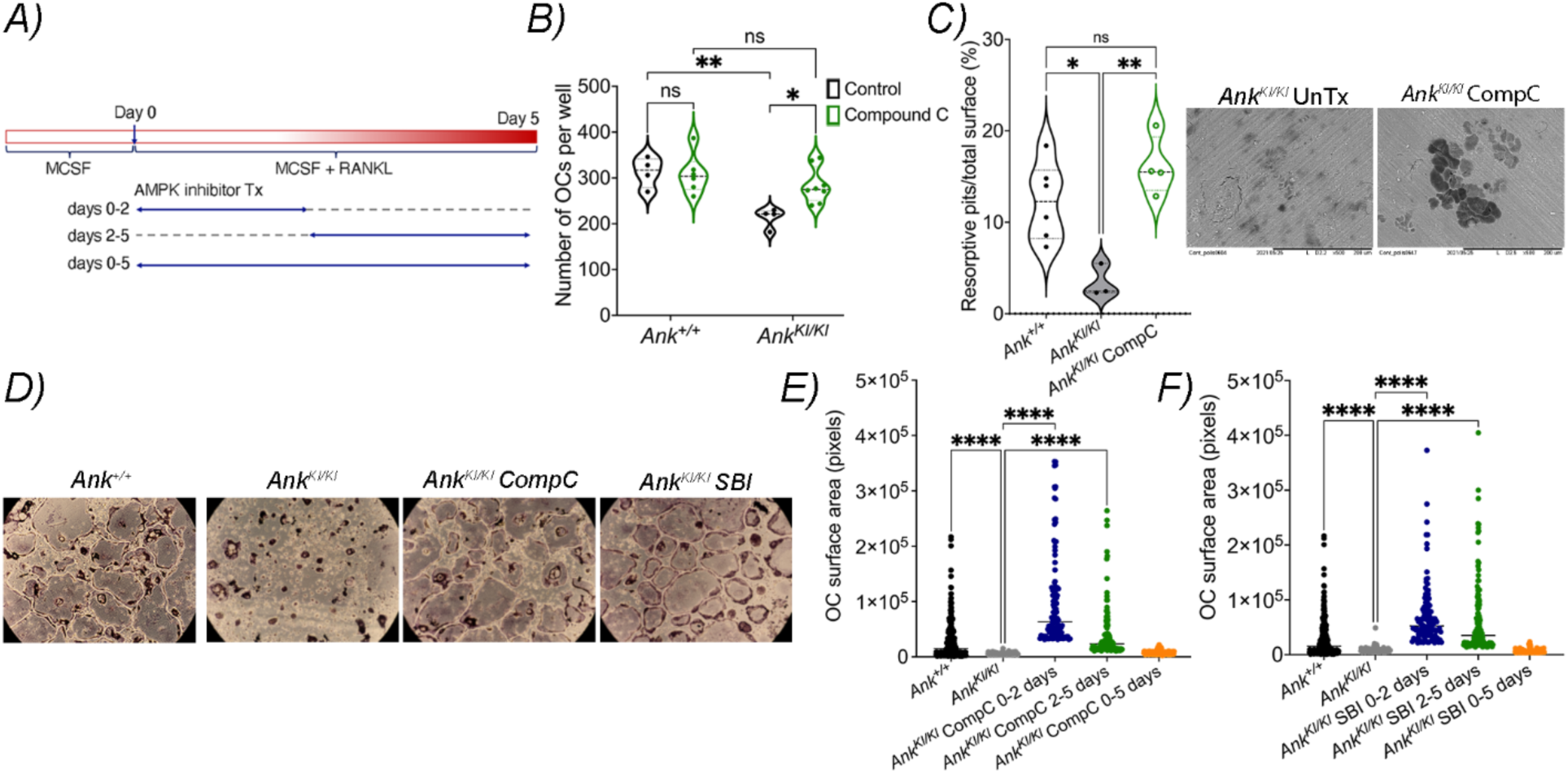
p-AMPK inhibitors compound C (CompC) and SBI-0206965 (SBI) rescue defects in *Ank^KI/KI^* BMM cultures. **A**) Schematic showing the schedule of AMPK inhibitor treatment during fusion stage (0-2 days), maturation stage (2-5 days), or continuous treatment (0-5 days) in BMM cultures. All groups were analyzed by the end of day 5 of M-CSF and RANKL treatment; **B**) Compound C treatment during fusion stage significantly increased *Ank^KI/KI^* OC numbers. Statistics were performed by two-way ANOVA followed by Tukey’s post hoc test; **C**) Compound C treatment during fusion stage significantly increased resorption pits in *Ank^KI/KI^* cultures shown by box and violin plot (left panel) and SEM images (right panel). Statistics were performed by one-way ANOVA followed by Tukey’s post hoc test; **D**) TRAP staining showed that CompC or SBI both restored small-sized OCs in *Ank^KI/KI^* cultures; **E**) CompC and **F**) SBI treatment during fusion or maturation stages both corrected small-sized *Ank^KI/KI^* OCs whereas continuous treatment had no rescue effects. Surface areas were compared to the *Ank^KI/KI^* untreated group. Bars indicate median values. Statistics were performed by one-way ANOVA followed by Tukey’s post hoc test to compare to *Ank^KI/KI^* no treatment group (\**p*<0.05, \*\**p*<0.01, \*\*\*\**p*<0.0001).

Without treatment, *Ank^KI/KI^* BMM cultures form reduced numbers of OCs that are smaller in size and have diminished resorption compared to *Ank^+/+^* BMMs.^(10)^ *Ank^KI/KI^* BMM cultures that were treated with compound C during days 0-2 showed restoration of OC numbers at day 5 (Figure 4B). More importantly, the compound C treatment on bone chip cultures from days 0-4 (fusion stage) also enhanced the bone resorption capability of *Ank^KI/KI^* BMMs analyzed 10 days after culture plating (Figure 4C). The restoration of osteoclastogenesis in *Ank^KI/KI^* cultures by compound C or SBI-0206965 corresponds to enlargement of OC size as shown in Figure 4D. Enlargement of *Ank^KI/KI^* OCs by AMPK inhibitors occurred either with treatment during days 0-2 or 2-5 while continuous treatment did not restore the *Ank^KI/KI^* OC size and was accompanied by reduced numbers of viable cells (Figure 4E & 4F; data not shown). These data suggest that increased p-AMPK has a negative effect on OC formation and bone resorption of *Ank^KI/KI^*BMMs *in vitro*. Treatment with p-AMPK inhibitors during the fusion stage successfully rescues the phenotype of *Ank^KI/KI^* OCs.

### AMPK inhibitors restore actin cytoskeleton and increase cellular acidosis

We suspected that the rescue effects of AMPK inhibitors are mediated via the restoration of cytoskeletal proteins that control the cell shape, size, and migration. Cytoskeletal proteins include actin, microtubules and intermediate filaments. Actin has been shown to be disrupted in *Ank^KI/KI^*OCs.^(10)^ We examined the effects of AMPK inhibitors on the actin cytoskeleton in *Ank^KI/KI^* OCs cultured on glass slides and bone chips. OCs cultured on glass slides typically formed one large actin ring whereas OCs cultured on bone chips presented multiple actin rings within one cell. *Ank^KI/KI^* OCs grown on glass slides had disrupted and smaller actin rings (Figure 5A). With compound C and SBI-0206965 treatment of *Ank^KI/KI^* OCs, the actin ring sizes were significantly enlarged (Figure 5A). *Ank^KI/KI^* OCs grown on bone chips had reduced actin ring sizes compared to *Ank^+/+^* OCs (top panels in Figure 5B). AMPK inhibitor treatment significantly restored the size of actin rings in *Ank^KI/KI^* cultures on bone chips (bottom panels in Figure 5B). These data suggest that inhibition of AMPK during the fusion stage restores the function of *Ank^KI/KI^* OCs partly by correcting actin dynamics.

**Figure 5.**
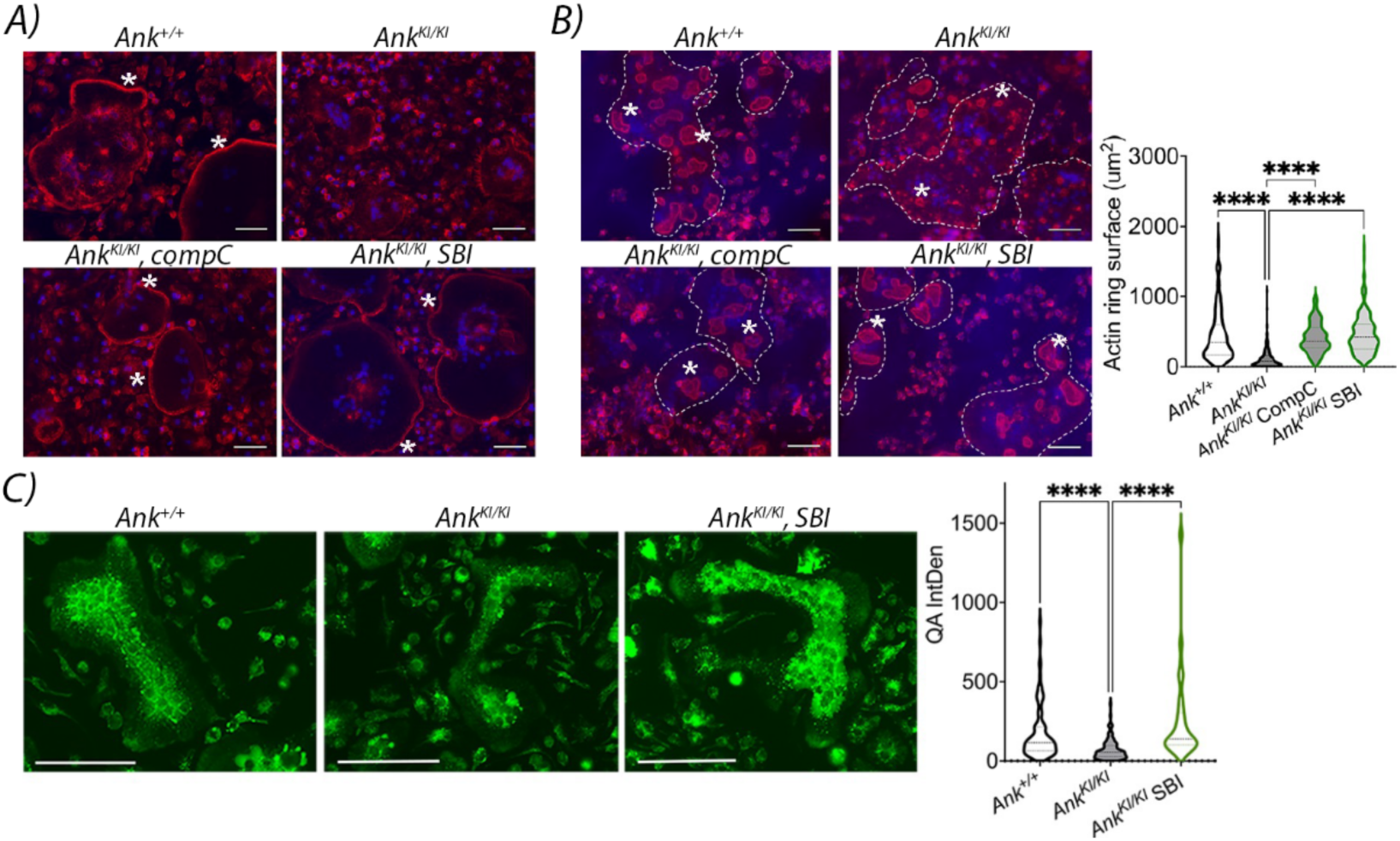
Rescue effects of AMPK inhibitors on actin cytoskeleton and cellular acidosis. **A**) *Ank^++^* and *Ank^KI/KI^* OCs cultured on glass slides stained by rhodamine phalloidin. Compound C (compC) and SBI-0206965 (SBI) treatment resulted in restoration of actin ring structure (*asterisks) in *Ank^KI/KI^* OCs; **B**) *Ank^++^* and *Ank^KI/KI^* OCs cultured on bone chips stained by rhodamine phalloidin. Dashed lines encircle one OC. Asterisks indicated actin rings on bone chips. Treatment of compC and SBI results in enlarged actin ring structures in *Ank^KI/KI^* cultures. Nuclei stained by Hoechst dye 33258. Scale bar = 50 μm; **C**) SBI treatment restored the reduced acidic vesicles in *Ank^KI/KI^* OCs stained by QA (green fluorescence). Scale bar = 100 μm. Box and violin plots in **B**) and **C**) indicate statistical significance by one-way ANOVA followed by Tukey’s post hoc test compared to *Ank^KI/KI^*untreated group. (\*\*\*\**p*<0.0001).

Acidosis plays an important role in osteoclastic bone resorption. We examined whether AMPK inhibitors act on acidic vesicles in OCs by quinacrine dihydrochloride (QA) dye. The protonated QA molecules are trapped inside the acidic vesicles and can be detected by fluorescence microscopy. We examined vesicle acidification in *Ank^+/+^* and *Ank^KI/KI^* OCs and found that there were less acidic vesicles in *Ank^KI/KI^* OCs shown by reduced QA intensities (Figure 5C). SBI-0206965 treatment restored the acidity in *Ank^KI/KI^* OCs (Figure 5C).

### Effects of AMPK inhibition on the skeletal phenotype of Ank^KI/KI^ mice

To study the effects of AMPK inhibition *in vivo,* we performed a pilot study to treat male and female *Ank^KI/KI^* mice with an AMPK inhibitor, SBI-0206965 (10 μg/g, once a week by IP injection), or vehicle for 7 weeks between age 1 to 8 weeks. *Ank^KI/KI^* mice weighed significantly less when compared to *Ank^+/+^* mice and SBI-0206965 treatment did not improve body weight gain in *Ank^KI/KI^* mice (data not shown). SBI-0206965 treatment did not correct the hyperostotic mandibles, trabecular and cortical parameters of femurs in *Ank^KI/KI^* mice (Figure 6A). Many μCT parameters including mandibular bone volume (BV), total volume (TV), femoral BV/TV, trabecular number (Tb.N), trabecular thickness (Tb.Th), and cortical thickness (Ct.Th) showed no significant differences between *Ank^KI/KI^* mice with vehicle and SBI-0206965 treatment (Table 1). *Ank^KI/KI^* mice that received SBI-0206965 injection have a tendency of increased periosteal and endocortical perimeters as well as cortical porosity (Table 1).

**Figure 6.**
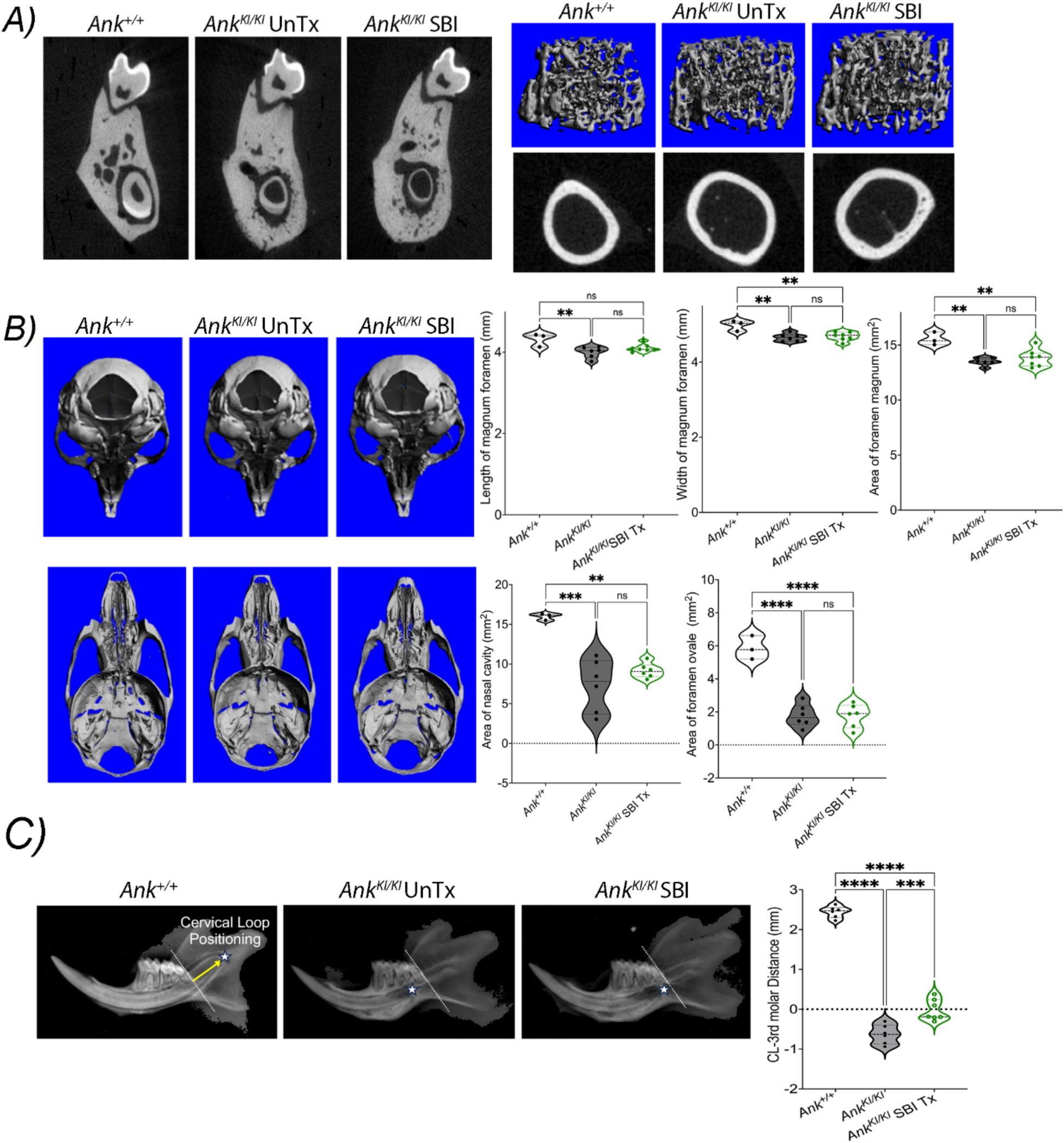
Representative μCT images of **A**) Mandibles, trabecular, and cortical bones; **B**) Skulls and cranial base. Histograms showing quantification of foramen magnum length, width, and area, the open space of nasal cavity, and the cranial foramina; **C**) Representative radiographs of mandibles and measurements of distance between cervical loop (CL, white *) and distal surface of 3^rd^ molar (dashed line) in *Ank^+/+^* mice and *Ank^KI/KI^* mice that received vehicle (UnTx) or SBI (SBI-0206965) injection. Statistics were performed by one-way ANOVA followed by Tukey’s post hoc test (\*\**p*<0.01, \*\*\**p*<0.0005, \*\*\*\**p*<0.0001).

**Table 1:** μCT measurements of mandibles and femurs. Data presented: Mean (standard deviation). TV: total volume; BV: bone volume; BV/TV: bone volume/total volume; Tb.N: trabecular number; Tb.Th: trabecular thickness; Tb.Sp: trabecular spacing; Tt.Ar: total area; Ct.Ar: cortical area; Ct.Th: cortical thickness; Ps.Pm: periosteal perimeter; Ec.Pm: endocortical perimeter; Porosity: cortical porosity. Statistics difference compared to *Ank^+/+^* group (*p<0.05) analyzed by one-way ANOVA followed by Tukey’s post hoc test.

| Mand. | *Ank*<sup>+/+</sup> | *Ank*<sup>KI/KI</sup> | *Ank*<sup>KI/KI</sup> SBI-0206965 |
| --- | --- | --- | --- |
| TV (mm<sup>3</sup>) | 2.66 (0.05) | 4.15 (0.34)* | 4.71 (0.61)* |
| BV (mm<sup>3</sup>) | 2.19 (0.05) | 3.42 (0.26)* | 3.87 (0.61)* |
| **Femoral trabecular bone** |  |  |  |
| BV/TV (%) | 11.49 (2.52) | 9.54 (1.48) | 11.37 (4.03) |
| Tb.N (1/mm) | 4.1 (0.46) | 3.9 (0.45) | 4.04 (0.69) |
| Tb.Th (mm) | 52.96 (1.1) | 50.76 (1.2) | 53.97 (1.8) |
| Tb.Sp (mm) | 246 (26.4) | 256 (30.8) | 249 (43.9) |
| **Femoral cortical bone** |  |  |  |
| Tt.Ar (mm<sup>2</sup>) | 1.35 (0.17) | 1.74 (0.27) | 1.79 (0.34) |
| Ct.Ar (mm<sup>2</sup>) | 0.6 (0.09) | 0.64 (0.07) | 0.69 (0.12) |
| Ct.Th (mm) | 0.16 (0.01) | 0.15 (0.006) | 0.15 (0.01) |
| Ps.Pm (mm) | 4.11 (0.26) | 4.67 (0.38) | 4.79 (0.49)* |
| Ec.Pm (mm) | 3.07 (0.17) | 3.72 (0.34)* | 3.76 (0.39)* |
| Porosity (%) | 0.27 (0.03) | 0.31 (0.0005) | 0.36 (0.0008) |

As seen in patients with CMD, *Ank^KI/KI^* mice have narrowed cranial foramina including the foramen magnum.^(7)^ *Ank^KI/KI^* mice administered with SBI-0206965 did not significantly enlarge foramen magnum, nasal cavity, and cranial foramina in comparison to *Ank^KI/KI^* mice treated with vehicle (Figure 6B). We previously showed that the cervical loop positioning of incisors in *Ank^KI/KI^* mice was significantly more ventrally located (before 3^rd^ molars) than in *Ank^+/+^* mice.^(28)^ This phenotype had been replicated in *Ank^+/+^* mice when bisphosphonate was injected suggesting that OC action controls the positioning of cervical loops of incisors. *Ank^KI/KI^* mice injected with SBI-0206965 had their cervical loops of incisors significantly more dorsally located than *Ank^KI/KI^* vehicle control mice (Figure 6C). These data suggest that systemic administration of SBI-0206965 under this regimen does not restore most of the CMD-like features but only partially rescues the localization of incisor cervical loops of *Ank^KI/KI^* mice.

## DISCUSSION

Several mechanisms of ATP efflux have been reported, including channel-mediated release, transporter-dependent release, exocytosis, and cell lysis.^(29)^ ANK loss-of-function results in reduced ATP efflux.^(14–16)^ Our data showed that ATP efflux was significantly reduced in *Ank^KI/KI^* BMMs, implicating reduced transporting activity of Ank_F377del_ protein. This is consistent with our previous findings that ANKH_F377del_/ANK_F377del_ identified in AD CMD showed reduced expression and function due to rapid degradation and mislocalization.^(11, 30)^ The AR form of CMD is caused by a R239Q mutation in Connexin 43 leading to mislocalization and reduced function of the Cx43 protein.^(31)^ Downregulation of Cx43 using siRNA prevents ATP efflux.^(32)^ We speculate that ATP homeostasis may be commonly affected in both the AD and AR forms of CMD.

Our initial assumption has been that reduction of cellular ATP release in *Ank^KI/KI^* BMMs will lead to accumulation of intracellular ATP. In contrast, we observed decreased intracellular ATP and ADP levels in *Ank^KI/KI^* BMMs. Interestingly, knockdown of Cx43 by the CRISPR-Cas9 method also leads to decreased intracellular ATP generation, due to impairment of mitochondrial function.^(33)^ ANK/ANKH and Cx43, two mutant proteins involved in CMD, both appear to mediate ATP efflux as well as intracellular ATP production. ATP generation mainly involves glycolysis, citric acid cycle, and oxidative phosphorylation.^(34)^ Glycolysis converts glucose into pyruvate. Subsequently, pyruvate is converted to acetyl coenzyme A which enters the citric acid cycle to produce one molecule of ATP, three molecules of NADH and one molecule of FADH2. Products from the citric acid cycle provide electrons for respiratory chain and oxidative phosphorylation, which produce the largest amounts of ATP. We showed significantly reduced plasma citric acid (isocitrate) in *Ank^KI/KI^* mice. Enrichment and pathway analyses further suggested the involvement of the citric acid cycle. Our data indicate that in addition to reduced transporting activity, the reduced ATP efflux and plasma citric acid in *Ank^KI/KI^* mice may result from impaired ATP metabolism inside cells.

Our metabolomics study identified potential plasma biomarkers for CMD in that citric acid and N-acetylneuraminic acid were significantly reduced in *Ank^KI/KI^* mice. Citric acid is the first step in the citric acid cycle when acetyl-CoA condenses with oxaloacetate. Reduced citric acid has been associated with several diseases, including rheumatoid arthritis, chronic kidney diseases, bipolar disorder, systemic lupus erythematosus and others.^(35–38)^ N-acetylneuraminic acid is the most common form of sialic acid, primarily concentrated in the central nervous system where it combines with gangliosides and glycoproteins.^(39)^ Sialic acid is known to interact with bacteria and viruses to affect their infectiveness and is linked to various inflammation-related diseases.^(40, 41)^ Although these two molecules are unlikely specific for CMD, these data suggest potential pathways affected by CMD-mutant ANK for future studies.

Extracellular and intracellular ATP acts directly as a signaling molecule via purinergic receptors or indirectly on AMPK.^(42, 43)^ Modulating ATP concentrations to rescue defective OCs in *Ank^KI/KI^* mice and CMD patients will be challenging because it was shown that ATP has biphasic effects on OCs. Treating OC cultures with low concentrations of ATP enhances OC formation and resorption whereas high concentrations of ATP results in cytotoxic and suppressive effects.^(42)^ AMPK is activated by low ATP levels and clinical trials using activators or inhibitors to regulate AMPK activity are ongoing. Our data showed that p-AMPK expression is increased during the fusion stage of *Ank^KI/KI^* OCs and inhibition of AMPK on fusing OCs restores the function of mutant OCs. These findings are in line with previous studies showing that AMPK is a negative regulator of osteoclastogenesis.^(22)^, ^(44, 45)^ The effects of AMPK on OBs are less consistent. *Prkaa1^−/−^* mice have increased OB-mediated bone formation whereas deletion of AMPK in OBs by crossing *Ampk*^−/−^ mice with Osx-cre mice showed significant reduction in trabecular bone volume.^(22),(46)^ Our future studies will be directed to other relevant skeletal cell types such as osteoblasts, osteocytes, and chondrocytes.

The interaction of actin with nucleotides has long been known. In 1950 it was discovered that ATP is an essential part of actin and that ATP is hydrolyzed to ADP and Pi upon actin polymerization.^(47, 48)^ Disturbed ATP homeostasis may at least in part explain the abnormal actin structure in *Ank^KI/KI^*OCs. AMPK has been shown to regulate the cytoskeleton by acting on integrin and cofilin.^(49–51)^ We examined levels of αϖβ3 integrin, which plays important roles in OC biology, including cell attachment, apoptosis, and resorption.^(52)^ AMPK inhibitors did not significantly alter αϖβ3 integrin expression and p-cofilin/cofilin levels in fusing *Ank^+/+^* and *Ank^KI/KI^* OCs or in mature OCs (data not shown). How AMPK inhibitors regulate actin-related molecular mechanisms in *Ank^KI/KI^* OCs remains to be determined. We used QA to track acidic vesicles in OCs because it is the more stable dye compared to acridine orange and LysoTracker.^(23)^ Another application of QA is for detecting intracellular ATP-containing vesicles.^(53)^ While the accumulation of QA into acidic vesicles is consistent among studies, the use of QA for detecting vesicular ATP storage is controversial.^(53, 54)^ One of the regulatory mechanisms for cellular acidity is the v-ATPase-AMPK axis.^(55–57)^ Increased AMPK activity inhibits the trafficking of v-ATPase, which decreases v-ATPase-mediated proton secretion.^(55, 56)^ The AMPK-dependent effect on the regulation of acid secretion is via direct phosphorylation of the v-ATPase V1A subunit affecting the distribution of cytosolic and membranous v-ATPase.^(58)^ Although commercial antibodies against mouse V0 and V1 subunits of v-ATPase do not show expression differences (data not shown), we speculate that the v-ATPase may be affected in *Ank^KI/KI^* mice via interference with V0V1 assembly/disassembly or its localization in mice.

High doses of SBI-0206965 (50 μg/g every 3 days for 30 days) has been injected to suppress cancer growth.^(59)^ At this concentration, SBI-0206965 also leads to impaired autophagy.^(59)^ Lower doses of SBI-0206965 (weekly injection of 10 μg/g at age of 15 weeks for 4 weeks) leads to increased numbers of F4/80-positive macrophages and decreased amounts of CD11b^+^Gr-1^+^ myeloid-derived suppressor cells (MDSCs).^(60)^ For *in vivo* experiments, we chose SBI-0206965 because of its high specificity and potency to inhibit AMPK activity compared to compound C.^(27)^ We opted for the lower dosage of SBI-0206965 (10 μg/g) to maximize effects on myeloid-derived OCs. Our *in vivo* data show partial rescue of the positioning of cervical loops of incisors, which is an OC-dependent phenotype.^(28)^ Failure to rescue other CMD-like features under this regimen may be due to 1) SBI-0206965 having undesired off-target effects. Both high (50 μg/g) and low doses (10 μg/g) SBI-0206965 injection result in impairment of autophagy and increased apoptosis;^(59, 60)^ 2) systemic administration of SBI-0206965 acting on many cell types, not just OCs. In addition to cell autonomous effects of AMPK on bone cells, AMPK may also exert effects on systemic pro-inflammatory cytokines;^(61, 62)^ 3) the inability to inhibit AMPK in *Ank^KI/KI^* mice only during the fusion stage of OCs. Our *in vitro* data showed that continuous inhibition of AMPK results in a worse OC phenotype in *Ank^KI/KI^* BMM cultures due to reduced cell viability. Our data of reduced cell viability are consistent with a previous study showing that compound C treatment increased cell death in MCF7 breast carcinoma cells.^(63)^

In summary, our study reveals a previously unrecognized role of energy metabolism in the pathogenesis of CMD in mice. We began to identify biomarkers for CMD by a metabolomics approach and showed an association of increased AMPK activation with impaired OC function in *Ank^KI/KI^* mice. This proof-of-principle study provides new insights into CMD pathogenesis and suggests several new directions for future CMD research.

## Author contribution

Ayano Hatori: Data curation; formal analysis; methodology; writing – review and editing. Shyam Kishor Sah: Data curation; formal analysis; methodology; writing – original draft; writing – review and editing. Koen van de Wetering: writing – review & editing. Ernst J Reichenberger: Conceptualization; supervision; writing – review & editing. I-Ping Chen: Conceptualization; data curation; formal analysis; funding acquisition; methodology; supervision; writing – original draft; writing – review & editing.

## Funding support

This work is the result of NIH funding (NIDCR DE025664 to IPC, NIAMS R01AR082460 and R21AR083597 to KvdW, and AR082440 to ERJ) and is subject to the NIH Public Access Policy. KvdW receives funding from PXE International.

## Acknowledgments

We appreciate the valued assistance from Dr. Jeremy Balsbaugh (Facility Director of Proteomic & Metabolomics Facility) and Dr. Timothy E. Moore (Director of Statistical Consulting Service).

## Conflict of Interest

The authors have declared that no conflict of interest exists.

## Ethics Statement

All experiments involving animals were performed in accordance with animal protocol AP-200644-0225 approved by the Animal Care Committee at the University of Connecticut Health (UConn Health).

## Data Availability Statement

Raw data of the current study are available from the corresponding author upon reasonable request.

## Notes

### Competing Interest Statement

The authors have declared no competing interest.

