## Supplemental file for "Disturbed ATP and AMPK homeostasis in an Ank_F377del_ mouse model for craniometaphyseal dysplasia"

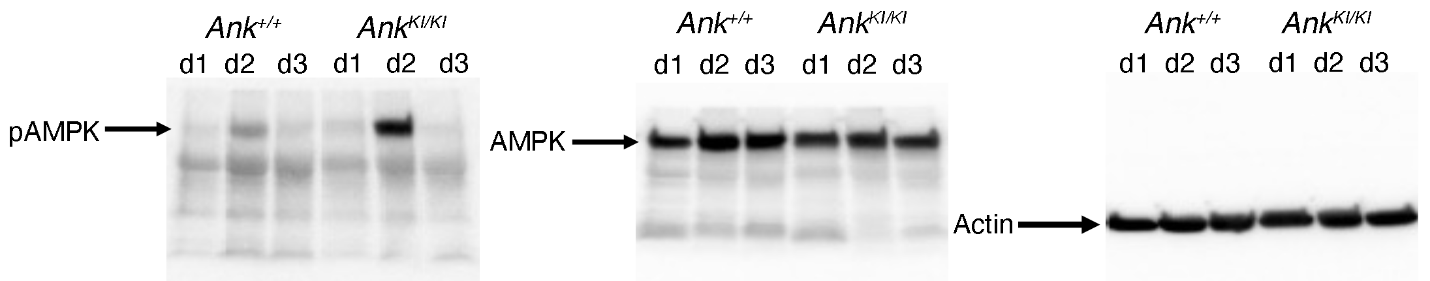

**Supplementary Figure S1.** Whole immunoblots of data presented in Figure 3.

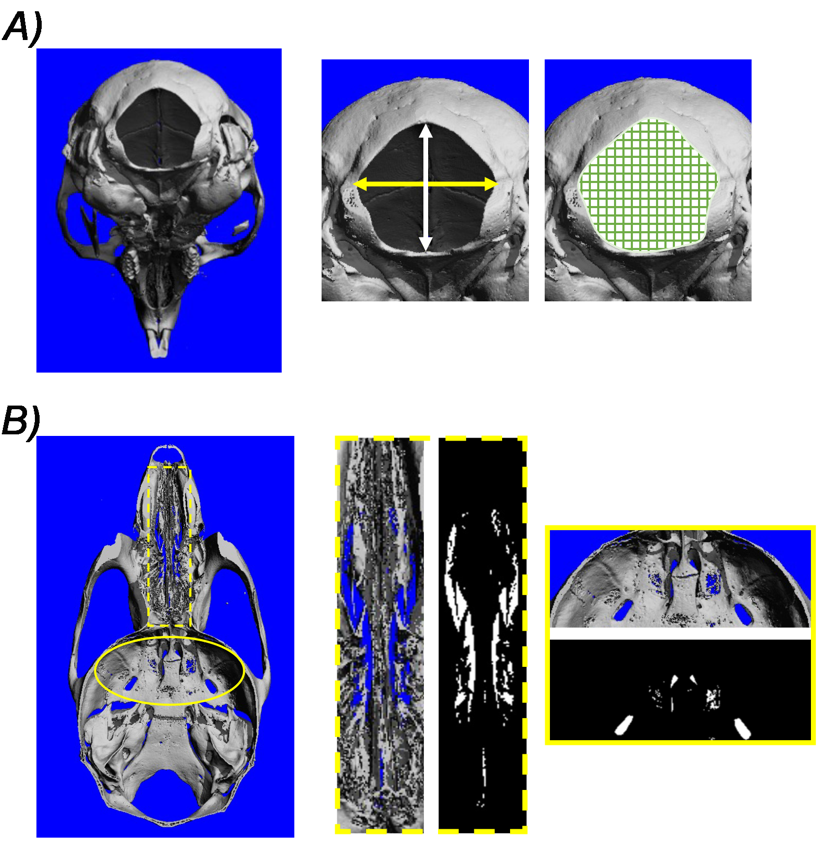

**Supplementary Figure S2:** μCT measurement of **A**) foramen magnum; white arrow indicating length, yellow arrow indicating width, and green grid surface showing area of foramen magnum; **B**) Open space (white area) of nasal cavity and cranial foramina using Image J software.

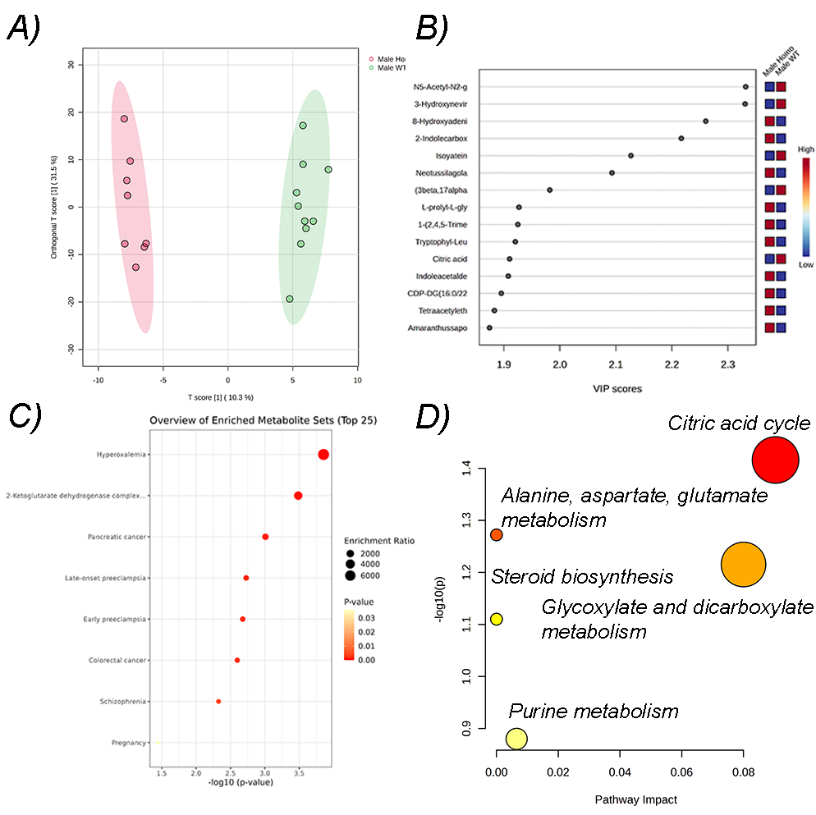

**Supplementary Figure S3**: Comparison of plasma metabolome in male *Ank^+/+^* and *Ank^KI/KI^* mice. **A**) PLS-DA of male data and **B**) VIP scores of top 15 differentiated metabolites; **C**) Enrichment assay showing enrichment ratio with *p*-value of upregulated and downregulated top 25 metabolites; **D**) Metabolic pathway analysis showing pathway impact and *p*-value of significantly upregulated and downregulated metabolites. (male *Ank^+/+^* n=10, male *Ank^KI/KI^* n=8).

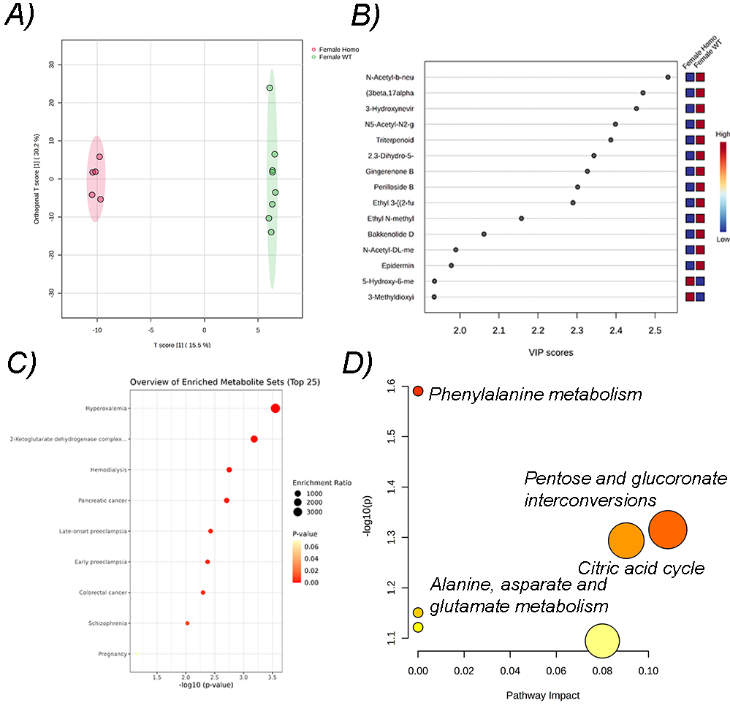

**Supplementary Figure S4**: Comparison of plasma metabolome in female *Ank^+/+^* and *Ank^KI/KI^* mice. **A**) PLS-DA of female data and **B**) VIP scores of top 15 differentiated metabolites; **C**) Enrichment assay showing enrichment ratio with *p*-value of upregulated and downregulated top 25 metabolites; **D**) Metabolic pathway analysis showing pathway impact and *p*-value of significantly upregulated and downregulated metabolites. (female *Ank^+/+^* n=8, female *Ank^KI/KI^* n=5).
